# Making predictions using poorly identified mathematical models

**DOI:** 10.1101/2023.11.26.568740

**Authors:** Matthew J. Simpson, Oliver J. Maclaren

## Abstract

Many commonly used mathematical models in the field of mathematical biology involve challenges of parameter non-identifiability. Practical non-identifiability, where the quality and quantity of data does not provide sufficiently precise parameter estimates is often encountered, even with relatively simple models. In particular, the situation where some parameters are identifiable and others are not is often encountered. In this work we apply a recent likelihood-based workflow, called *Profile-Wise Analysis* (PWA), to non-identifiable models for the first time. The PWA workflow addresses identifiability, parameter estimation, and prediction in a unified framework that is simple to implement and interpret. Previous implementations of the workflow have dealt with idealised identifiable problems only. In this study we illustrate how the PWA workflow can be applied to both structurally non-identifiable and practically non-identifiable models in the context of simple population growth models. Dealing with simple mathematical models allows us to present the PWA workflow in a didactic, self–contained document that can be studied together with relatively straightforward Julia code provided on GitHub. Working with simple mathematical models allows the PWA workflow prediction intervals to be compared with *gold standard* full likelihood prediction intervals. Together, our examples illustrate how the PWA workflow provides us with a systematic way of dealing with non-identifiability, especially compared to other approaches, such as seeking ad hoc parameter combinations, or simply setting parameter values to some arbitrary default value. Importantly, we show that the PWA workflow provides insight into the commonly–encountered situation where some parameters are identifiable and others are not, allowing us to explore how uncertainty in some parameters, and combinations of some parameters, regardless of their identifiability status, influences model predictions in a way that is insightful and interpretable.

## 1 Introduction

Interpreting experimental data using mechanistic mathematical models provides an objective foundation for discovery, prediction, and decision making across all areas of science and engineering. Interpreting data in this way is particularly relevant to applications in the life sciences where new types of measurements and data are continually being developed. Therefore, developing computational methods that can be used to explore the interplay between experimental data, mathematical models, and real-world predictions is of broad interest across many application areas in the life sciences.

Key steps in using mechanistic mathematical models to interpret data include: (i) identifiability analysis; (ii) parameter estimation; and, (iii) model prediction. Recently, we developed a systematic, computationally-efficient workflow, called *Profile-Wise Analysis* (PWA), that addresses all three steps in a unified way [63]. In essence, the PWA workflow involves propagating profile-likelihood-based confidence sets for model parameters, to model prediction sets by isolating how different parameters, and different combinations of parameters, influences model predictions [63]. This enables us to explore how variability in individual parameters, and variability in groups of parameters directly influence model predictions in a framework that is straightforward to implement and interpret in terms of the underlying mechanisms encoded into the mathematical models [62, 63]. This insight is possible because we harness the *targeting* property of the profile likelihood to isolate the influence of individual parameters [14,21,23,50,51], and we propagate forward from parameter confidence sets into prediction confidence sets. This feature is convenient since we can target individual interest parameters, or lower-dimensional combinations of parameters, one at-a-time, thereby providing mechanistic insight into the parameter(s) of interest, as well as making our approach scalable since we can deal with single parameters, or combinations of parameters, without needing to evaluate the entire likelihood function, only *profiles* of it. Profile-wise prediction confidence sets can be combined in a very straightforward way to give an overall curvewise prediction confidence set that accurately approximates the gold standard full likelihood prediction confidence set. One of the advantages of the PWA workflow is that it naturally leads to curvewise prediction sets that avoids challenges of computational overhead and interpretation that are inherent in working with pointwise approaches [31, 39]. The first presentation of the PWA work-flow focused on idealised identifiable models only, where the quality and quantity of the available data meant that all parameters were identifiable. Here we address to the more practical scenario of exploring how the PWA workflow applies to non-identifiable models. This is important because non-identifiability is routinely encountered in mathematical biology, and other modelling applications in the life sciences [27].

The first component of the PWA workflow is to assess parameter identifiability. Parameter identifiability is often considered in terms of *structural* or *practical* identifiability [79]. Structural identifiability deals with the idealised situation where we have access to an infinite amount of ideal, noise-free data [17, 18]. Structural identifiability for mathematical models based on ordinary differential equations (ODE) is often assessed using software that includes DAISY [8] and GenSSI [41]. In brief, GenSSI uses Lie derivatives of the ODE model to generate a system of input-output equations, and the solvability properties of this system provides information about global and local structural identifiability. In contrast, practical identifiability deals with the question of whether finite amounts of imperfect, noisy data is sufficient to identify model parameters [38, 52–54]. Practical identifiability can be implicitly assessed through the failure of sampling methods to converge [32, 58, 59]. Other approaches to determine practical identifiability include indirectly analysing the local curvature of the likelihood [11, 18, 30, 73] or directly assessing the global curvature properties of the likelihood function by working with the profile likelihood [52–54]. Therefore, practical identifiability is a more strict requirement than structural identifiability, as demonstrated by the fact that many structurally identifiable models are found to be practically non-identifiable when considering the reality of dealing with finite amounts of noisy data [59, 61]. Fröhlich et al. [27] compared several computational approaches (bootstrapping, profile likelihood, Fisher information matrix, and multi-start based approaches) to diagnose structural and practical identifiability of several models encountered in the systems biology literature and concluded that the profile likelihood produced reliable results, whereas the other approaches could be misleading.

In this work we will explore how the PWA workflow can be applied to both structurally non-identifiable and practically non-identifiable mathematical models that are routinely encountered in modelling population biology phenomena. Though we have not explicitly considered PWA and profile likelihood methods for non-identified models previously, there is reason to expect that it may perform reasonably well despite these challenges. As mentioned above, previous work in the context of systems biology by Fröhlich et al. [27] found that profile likelihood is the only method they consider that performs well in the presence of practical identifiability [27]. Investigations in other application areas has led to similar observations. In particular, in econometrics, Dufour [24] considers the construction of valid confidence sets and tests for econometric models under poor identifiability. They first show that, lacking other constraints, valid confidence sets must typically be unbounded. They then show that Wald-style confidence intervals of the form ‘estimate plus or minus some number of standard errors’ lack this property and are always bounded. In contrast, likelihood-based methods, such as likelihood ratios and associated confidence intervals, can produce appropriately unbounded intervals that maintain correct (or higher) asymptotic coverage. Zivot et al. [81] similarly compare confidence intervals for poorly identified econometric models based on inverting Lagrange multiplier, likelihood ratio, and Anderson-Rubin tests to Wald-style confidence intervals. As before, Zivot et al. [81] show that the relatively flat likelihood corresponds to appropriately wide confidence intervals, maintaining or exceeding coverage is maintained. Further, they demonstrate that likelihood-based intervals perform much better than Wald intervals [81]. In particular, these intervals may generally be unbounded unless additional constraints are applied, as shown to be theoretically necessary by Dufour [24]. Zivot et al.’s study is computational, while [24, 74] supplement this with additional asymptotic arguments. A more recent review of these and similar results, again in econometrics, is given by Andrews et al. [2], who also discuss how naive application of the bootstrap typically fails in the poorly identified setting (which is consistent with our recent results in [75] and those of [27]). In light of these results we expect that likelihood-based confidence intervals for parameters in poorly identified problems may still perform well. Though these will be unbounded in general, they will typically rule out some regions of the parameter space corresponding to e.g., rejected values of identified parameter combinations or one-sidedly bounded intervals. Furthermore, in most problems we can combine these intervals with simple *a priori* bounds (which are weaker assumptions than prior probability distributions; see [65]) to obtain confidence sets with the same coverage as the unbounded sets (again, see [65]).

In the context of prediction, our approach begins with a *gold-standard* full likelihood method that carries out prediction via initial parameter estimation in such a way that coverage for predictions is at least as good as coverage for parameters. Thus, the desirable properties for parameter confidence sets are expected to hold up for predictions, and predictions can also be thought of as functions or functionals of the parameters that may be better identified than the underlying parameters. The benefit of going via the parameter space rather than directly to predictions is that we can more easily generalise to new scenarios with mechanistic model components and parameters. In addition to this gold standard approach, we then introduce the PWA approach, which is based on the same general principles but requires less computation in exchange for potentially lower coverage. PWA also facilitates an understanding of the connection between individual parameters and predictions [63]. Guided by these theoretical intuitions about general models, we carry out computational experiments for particular models. One limitation of our approach is that the theoretical underpinnings are asymptotic and that finite sample corrections may be needed when data is sparse and the parameter space is large. One approach to address this would be applying the calibration methods from [75], which can be used with minimal modifications to our overall workflow. We leave this for future work.

An mentioned above, understanding the ability of the PWA workflow to deal with non-identifiable models is important because many mathematical models in the field of mathematical biology are non-identifiable. To demonstrate this point, we will briefly recall the history of mathematical models of avascular tumour spheroid growth. The study of tumour spheroids has been an active area of research for more than 50 years, starting with the seminal Greenspan model in 1972 [29]. Greenspan’s mathematical model involves a series of analytically tractable boundary value problems whose solutions provide a plausible explanation for the observed three–stage, spatially structured growth dynamics of avascular *in vitro* tumour spheroids [29]. Inspired by Greenspan, more than 50 years of further model development and refinement delivered continual generalisations of this modelling framework. This iterative process produced many detailed mathematical models that have incorporated increasingly sophisticated biological detail. This progression of ideas led to the development of time-dependent partial differential equation (PDE)-based models [13], moving boundary PDE models [12], multi-phase PDE models with moving boundaries [76, 77], and multi-component PDE models that simultaneously describe spheroid development while tracking cell cycle progression [35]. These developments in mathematical modelling sophistication did not consider parameter identifiability and it was not until very recently that questions of parameter identifiability of these models was assessed, finding that even the simplest Greenspan model from 1972 turns out to be practically non-identifiable when using standard experimental data focusing on the evolution of the outer spheroid radius only [11, 46]. This dramatic example points to the need for the mathematical biology community to consider parameter identifiability in parallel with model development to ensure that increased modelling capability and modelling detail does not come at the expense of those models to deliver practical benefits.

Previous approaches for dealing with non-identifiable models in mathematical and systems biology and related areas are to introduce some kind of simplification by, for example, setting parameters to default values [32, 59, 73], or seeking to find parameter combinations that simplify the model [21, 70]. For structurally non-identifiable models, symbolic methods can be used to find locally identifiable reparameterisations (e.g., [16,20], summarised in [21]), though these are difficult to apply to the practically non-identifiable case and typically require symbolic derivatives to be available. Several approaches can be used to find parameter combinations in the practically non-identifiable scenario, such as the concept of *sloppiness* [4, 5, 30] where a log-transformation is used to explore the possibility of finding informative combinations of parameters, typically in the form of ratios of parameters or ratios of powers of parameters as a means of simplifying the model. In this work, we take a different approach and use the (expected [24]) desirable properties of likelihood-based confidence intervals for poorly identified models, along with the the targeting property of the profile likelihood as used within the PWA workflow, to propagate confidence sets in parameter values to confidence sets for predictions for non-identifiable parameters. This approach provides insight because the targeting property of the profile likelihood can illustrate that some parameters are identifiable while others are not, and importantly the PWA workflow directly links the relevance of different parameters, regardless of their identifiability, to model predictions. Overall, we illustrate that the PWA workflow can be used as a systematic platform for making predictions, regardless of whether dealing with identifiable or non-identifiable models. This is useful for the commonly-encountered situation of *partly* identifiable models where some parameters are well–identified by the data and others are not [59]. To make this presentation as accessible as possible we present results for very simple, widely adopted mathematical models relating to population dynamics. Results are presented in terms of two simple case studies, the first involves structural non-identifiability, and the second involves practical non-identifiability. The first, structurally non-identifiable case is sufficiently straightforward that all calculations are performed by discretising the likelihood function and using appropriate grid searches to construct various profile likelihoods. This approach has the advantage of being both conceptually and computationally straightforward. The second, practically non-identifiable case, is dealt with using numerical optimisation instead of grid searches. This approach has the advantage of being computationally efficient and more broadly applicable than using simple grid searchers. As we show, the PWA leads to sensible results for these two simple case studies, and we anticipate that the PWA will also leads to useful insights for other non-identifiable models because related likelihood-based approaches are known to perform well in the face of non-identifiability [24, 27, 75, 81]. Open source software written in Julia is provided on GitHub to replicate all results in this study.

## 2 Results and Discussion

We present this these methods in terms of two case studies that are based on very familiar, mathematical models of population dynamics. Before presenting the PWA details, we first provide some background about these mathematical models and briefly discuss some application areas that motivate their use.

### 2.1 Logistic population dynamics

The logistic growth model

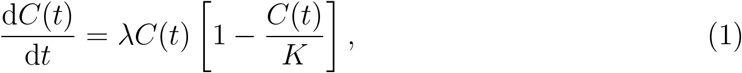

with solution

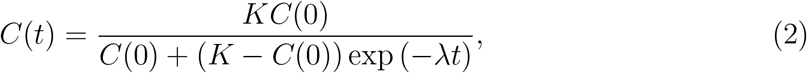

is one of the most widely-used mathematical models describing population dynamics [26, 37,49]. This model can be used to describe the time evolution of a population with density *C*(*t*) *>* 0, where the density evolves from some initial density *C*(0) to eventually reach some carrying capacity density, *K >* 0. For typical applications with *C*(0)*/K ≪* 1 the solution describes near exponential growth at early time, with growth rate *λ*, before the density eventually approaches the carrying capacity in the long time limit, *C*(*t*) *→ K*^*−*^ as *t → ∞*. One of the reasons that the logistic growth model is so widely employed is that it gives rise to a sigmoid shaped solution curve that is so often observed in a range of biological phenomena across a massive range of spatial and temporal scales. Images in Figure 1 show typical applications in population biology where sigmoid growth is observed. Figure 1(a) shows the location of a coral reef on the Great Barrier Reef that is located off the East coast of Australia. Populations of corals are detrimentally impacted by a range of events, such as tropical cyclones or coral bleaching associated with climate change [34]. Data in Figure 1(b) shows measurements of the percentage of total hard coral cover after an adverse event that reduced the proportion of area covered by hard coral to just a few percent [25,61,62]. The recovery in hard coral cover took more than a decade for the corals to regrow and colonise the reef, back to approximately 80% area occupancy. This recovery data can be described using the logistic growth model, where it is of particular interest to estimate the growth rate *λ* because this estimate can be used to provide a simple estimate of the timescale of regrowth, 1*/λ* [61, 62].

**Figure 1:**
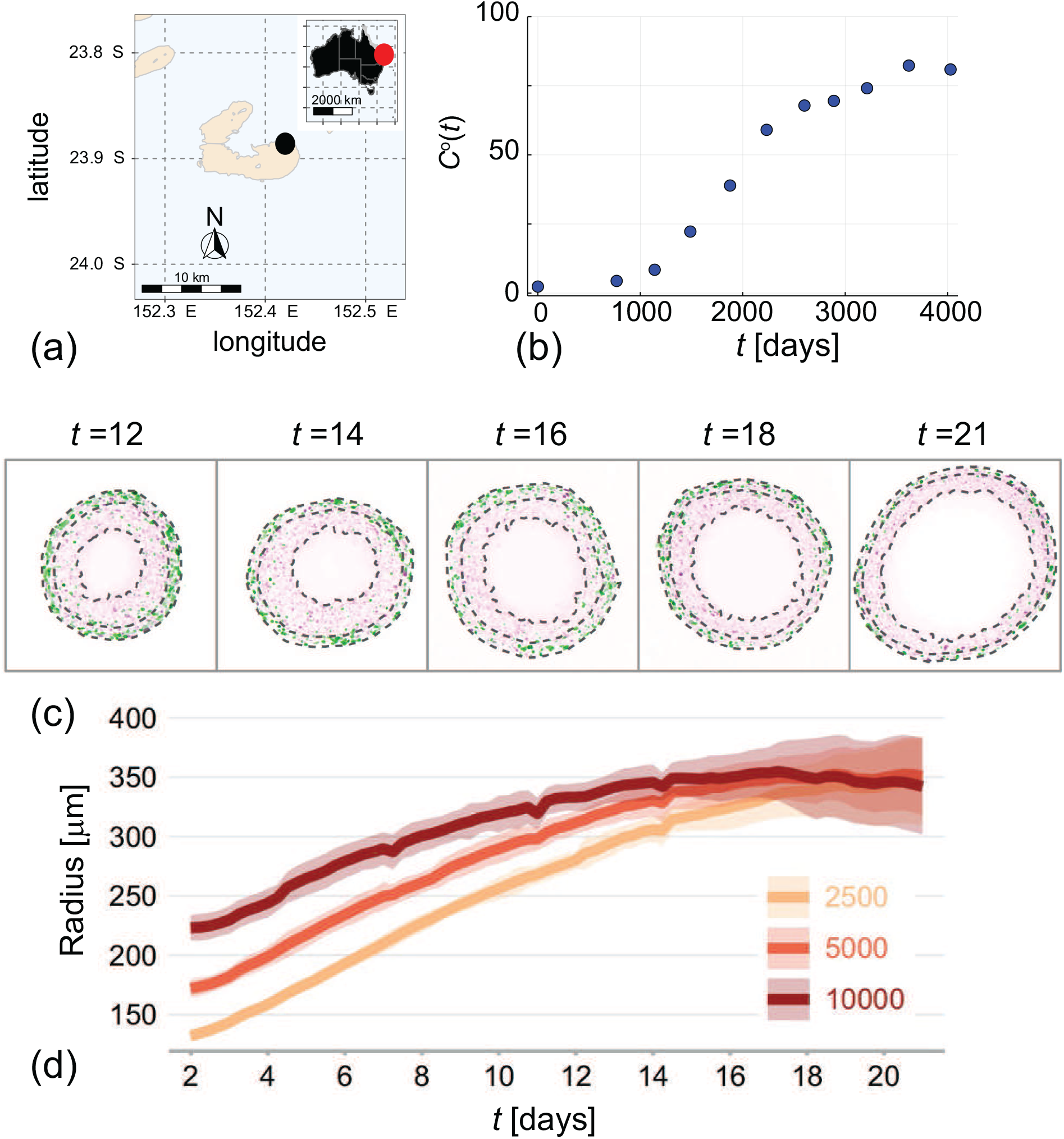
Practical applications of logistic growth models in ecology and cell biology. (a) Location of Lady Musgrave Island (black disc) relative to the Australian mainland (inset, red disc). (b) Field data showing the time evolution of the percentage total hard coral cover, *C*°(*t*) (blue discs) after some disturbance at Lady Musgrave Island monitoring site 1 [25]. Images in (a)–(c) reproduced with permission from Simpson et al. (2023) [62]. (c) Experimental images showing equatorial slices through tumour spheroids grown with melanoma cells. Images from left–to–right show spheroids harvested at 12, 14, 16, 18 and 21 days after formation [10]. Each cross section shows that the spheroids grow as a three– layer compound sphere with the inner–most sphere containing dead cells, the central spherical shell containing live but arrested cells that are unable to progress through the cell cycle, and the outer–most spherical shell contains live cells progressing through the cell cycle. (d) Summarises the dynamics of the outer most radius of a group of spheroids of the same melanoma cell line grown from different numbers of cells, including spheroids initiated with 2500, 5000 and 10000 cells, as indicated. Images in (c)–(d) reproduced with permission from Browning et al. (2021) [10].

Another classical application of the logistic growth model is to understand the dynamics of populations of tumour cells grown as *in vitro* avascular tumour spheroids. For example, the five experimental images in Figure 1(c) show slices through the equatorial plane of a set of *in vitro* melanoma tumour spheroids, after 12, 14, 16, 18 and 21 days of growth after spheroid formation [10, 47]. The structure of these spheroids approximately takes the form of a growing compound sphere. Dotted lines in Figure 1(c) show three boundaries: (i) the outer-most boundary that defines the outer radius of the spheroid; (ii) the central boundary that separates a cells according to their cell cycle status; and, (iii) the inner-most boundary that separates the central shell composed of living but non-proliferative cells from the central region that is composed of dead cells. Cells in the outer-most spherical shell have access to sufficient nutrients that these cells are able to proliferate, whereas cells in the central spherical shell have limited access to nutrients which prevents these cells entering the cell cycle [10]. Data in Figure 1(d) shows the time evolution of the outer radius for groups of spheroids that are initiated using different numbers of cells (i.e. 2500, 5000, 10000 cells, as indicated). Regardless of the initial size of the spheroids, we observe sigmoid growth with the eventual long–time spheroid radius is apparently independent of the initial size. This sigmoid growth behaviour is consistent with the logistic growth model, Equation 1, where here we consider the variable *C*(*t*) to represent the spheroid radius [47]. In addition to modelling coral reef re-growth [62] and tumour spheroid development [10, 11], the logistic model has been used to model a wide range of cell biology population dynamics including *in vitro* wound healing [42] and *in vivo* tumour dynamics [28, 57, 78]. Furthermore, the logistic equation and generalisations thereof are also routinely used in various ecological applications including modelling populations of sharks [33], polyps [44], trees [1] and humans [66].

In terms of mathematical modelling, the simplicity of the logistic growth model introduces both advantages and disadvantages. The advantages of the logistic growth model include the fact that it is both conceptually straightforward and analytically tractable. In contrast, the simplicity of the logistic growth model means that it is not a high–fidelity representation of biological detail. Instead, it would be more accurate to describe the logistic growth model as a phenomenologically–based model that can be used to provide insightful, but indirect information about biological population dynamics. In response to the disadvantages of the logistic model, there have been many generalisations proposed, such as those reviewed by Tsoularis and Wallace [69]. For the purposes of this study we will work with two commonly–used generalisations. As we explain later, these two commonly–used generalisations introduce different challenges in terms of parameter identifiability.

The first generalisation we will consider is to extend the logistic growth model to include a separate linear sink term,

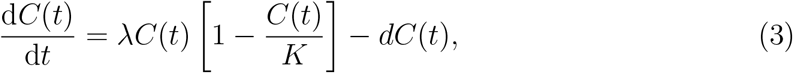

where *d >* 0 is a rate of removal associated with the linear sink term. Incorporating a linear sink term into the logistic growth model has been used to explicitly model population loss, either through some kind of intrinsic death process [6], a death/decay associated with external signals, such as chemotherapy in a model of tumour cell growth [68] or external harvesting process [9, 45], as is often invoked in models of fisheries [22, 80]. Re–writing Equation 3 as

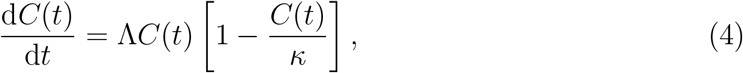

with Λ = *λ− d* and *κ* = *K*Λ*/λ*, with *λ >* 0, preserves the structure of the logistic growth equation so that the solution is

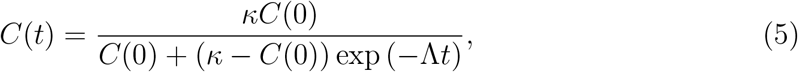

which now gives three long-term possibilities: (i) *C*(*t*) *→ K* as *t → ∞* if *λ > d*: (ii) *C*(*t*) *→* 0 as *t → ∞* if *λ < d*, and (iii) *C*(*t*) = *C*(0) for all 0 *< t < ∞* for the special case *λ* = *d*. In this work we will refer to Equation 3 as the *logistic model with harvesting* [9].

The second generalisation we will consider is to follow the work of Richards [55] who generalised the logistic growth model to

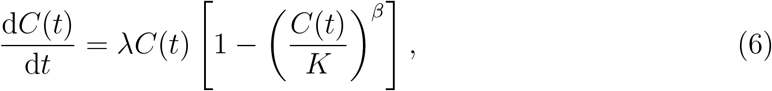

where the free parameter *β >* 0 was introduced by Richards to provide greater flexibility in describing a wider range of biological responses in the context of modelling stem growth in seedlings [55]. This model, which is sometimes called the *Richards’ model*, has since been used to study population dynamics in a range of applications, including breast cancer progression [67]. The solution of the Richards’ model can be written as

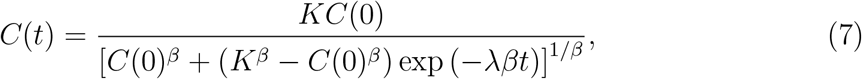

which clearly simplifies to the solution of the logistic equation when *β* = 1. The Richards’ model is also related to the well–known von Bertalanffy growth model in which *β* = 1*/*3 [69]. Instead of considering fixed values of *β*, the Richards’ model incorporates additional flexibility by treating *β* as a free parameter that can be estimated from experimental data.

This work focuses on the development and deployment of likelihood–based methods to make predictions with poorly–identified models. A preliminary observation about the mathematical models that we have presented so far is that both the logistic model, Equation 1, and the Richards’ model, Equation 6 are structurally identifiable in terms of making perfect, noise–free observations of *C*(*t*) [41]. In contrast, the logistic model with harvesting is structurally non-identifiable since making perfect, noise–free observations of *C*(*t*) enables the identification of Λ and *κ* only, for which there are infinitely many choices of *λ, d* and *K* that give the same *C*(*t*). We will now illustrate how the PWA workflow for parameter identifiability, estimation and prediction can be implemented in the face of both structural and practical non-identifiability.

### 2.2 Logistic model with harvesting

To solve Equation 3 for *C*(*t*) we must specify an initial condition, *C*(0), together with three parameters *θ* = (*λ, d, K*)^⊤^. We proceed by assuming that the observed data, denoted *C*°(*t*), corresponds to the solution of Equation 3 corrupted with additive Gaussian noise with zero mean and constant variance so that *C*°(*t*) | *θ* = *N* (*C*(*t*_*i*_), *σ*^2^), where *C*(*t*_*i*_) is the solution of Equation 3 at a series of *I* time points, *t*_*i*_ for *i* = 1, 2, 3, …, *I*. Within this standard framework the loglikelihood function can be written as

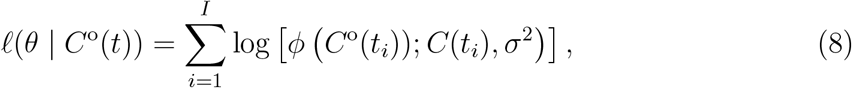

where *ϕ*(*x*; *µ, σ*^2^) denotes a Gaussian probability density function with mean *µ* and variance *σ*^2^, and *C*(*t*) is the solution of Equation 3. We treat *ℓ*(*θ* | *C*°(*t*)) as a function of *θ* for fixed data [15], and in this first example, for the sake of clarity and simplicity, we treat *C*(0) and *σ* as known quantities [32]. This has the benefit of reducing this problem to dealing with three unknown parameters, *θ* = (*λ, β, K*)^⊤^. This assumption will be relaxed later in Section 2.3. Synthetic data is generated with *λ* = 0.01, *d* = 0.002, *K* = 100, *C*(0) = 5 and *σ* = 5. Data is collected at *t*_*i*_ = 100 *×* (*i −* 1) for *i* = 1, 2, 3, …, 11 and plotted in Figure 2(a). The remainder of this mathematical exploration is independent of the units of these quantities, however to be consistent with the data in Figure 1(b) the dimensions of *C*(*t*) and *K* would be taken to be densities expressed in terms of a percentage of area occupied by growing hard corals, and the rates *λ* and *d* would have dimensions of [*/*day]. Similarly, work with the data in Figure 1(d) the variables *C*(*t*) and *K* would be taken to represent spheroid radius with dimensions *µ*m, and the rates *λ* and *d* would have dimensions [*/*day] [47].

**Figure 2:**
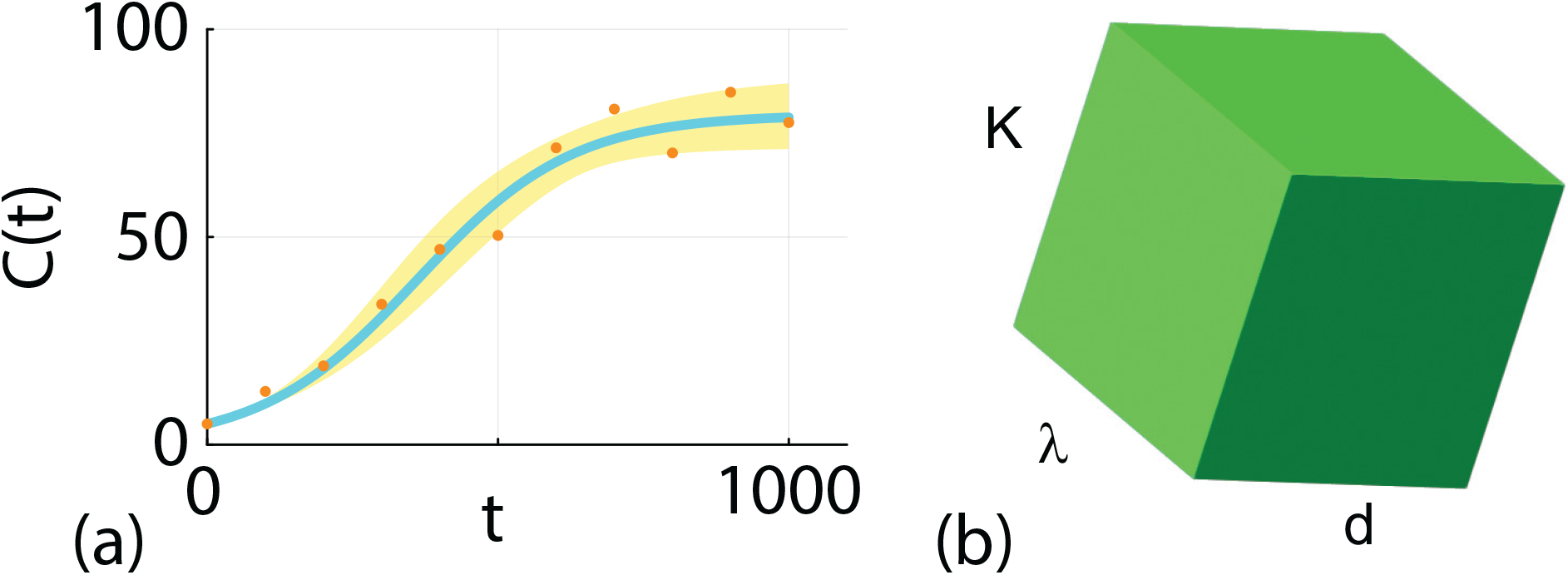
(a) Synthetic data (orange dots) generated by taking the solution of Equation 4 with *λ* = 0.01, *d* = 0.002, *K* = 100, *C*(0) = 5 at *t*_*i*_ = 100 *×* (*i −* 1) for *i* = 1, 2, 3, …, 11 and incorporating additive Gaussian noise with zero mean and *σ* = 5. The MLE solution (solid blue) corresponds to the solution of Equation 4 with 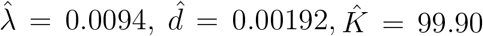 with *C*(0) = 5. The 95% prediction interval (shaded gold) is obtained by propagating parameter values where 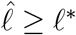 forward to construct the prediction envelope. Here *ℓ*^*∗*^ = *−*Δ_0.95,3_*/*2 = *−*3.91, where Δ_*q,n*_ refers to the *q*th quantile of the χ^2^ distribution with *n* degrees of freedom [56]. (b) Schematic of the (*λ, d, K*)^⊤^ parameter space.

We now apply the PWA framework for identifiability, estimation and prediction by evaluating *ℓ*(*θ* | *C*°(*t*)) across a broad region of parameter space that contains the true values: 0.0001 *< λ <* 0.05; 0 *< d <* 0.01; and 50 *< K <* 200. Since we are dealing with a relatively simple three–dimensional parameter space, shown schematically in Figure 2(b), we can explore different parameter combinations by taking a uniform 500^3^ regular discretisation of the parameter space and evaluate *ℓ*(*θ* | *C*°(*t*)) at each of the 500^3^ parameter combinations defined by the uniform discretisation. Calculating the maximum value of *ℓ*(*θ* | *C*°(*t*)) over the 500^3^ values, 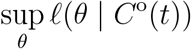, allows us to work with the normalized loglikelihood function

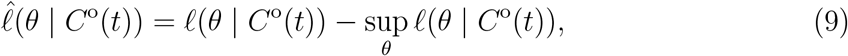

so that the maximum normalised loglikelihood is zero. The value of *θ* that maximises *ℓ*(*θ* | *C*°(*t*)) is denoted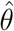, which is called the maximum likelihood estimate (MLE). In this instance we have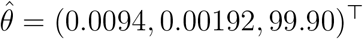, which is reasonably close to the true value, *θ* = (0.01, 0.002, 100)^⊤^, but this point estimate provides no insight into inferential precision which is associated with the curvature of the likelihood function [21, 50].

To assess the identifiability of each parameter we construct a series of univariate profile likelihood functions by partitioning the full parameter *θ* into interest parameters *ψ* and nuisance parameters *ω*, so that *θ* = (*ψ, ω*). For a set of data *C*°(*t*), the profile log-likelihood for the interest parameter *ψ*, given the partition (*ψ, ω*), is

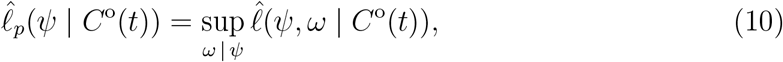

which implicitly defines a function *ω*^*∗*^ (*ψ*) of optimal values of *ω* for each value of *ψ*, and defines a surface with points (*ψ, ω*^*∗*^(*ψ*)) in parameter space. In the first instance we construct a series of univariate profiles by taking the interest parameter to be a single parameter, which means that (*ψ, ω*^*∗*^(*ψ*)) is a univariate curve that allows us to visualise the curvature of the loglikelihood function.

Since this first example deals with a relatively straightforward three–dimensional log-likelihood function where we have already evaluated 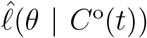 on a 500^3^ regular discretisation of *θ*, the calculation of the profile likelihood functions is straightforward. For example, if our interest parameter is *ψ* = *λ* and the associated nuisance parameter is *ω* = (*d, K*)^⊤^, for each value of *λ* along the uniform discretisation of the interval 0.0001 *< λ*_*j*_ *<* 0.05 for *j* = 1, 2, 3, …, 500, we compute the indices *k* and *l* that maximises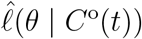, where *θ* is evaluated on the uniform discretisation (*λ*_*j*_, *d*_*k*_, *K*_*l*_). This *optimisation* calculation can be performed using a straightforward grid search. Repeating this procedure by setting *ψ* = *d* and *ω* = (*λ, K*)^⊤^, and *ψ* = *K* and *ω* = (*λ, d*)^⊤^ gives univariate profile likelihoods for *d* and *K*, respectively. The three univariate profile likelihood functions are given in Figure 3 where each profile likelihood function is superimposed with a vertical green line at the MLE and a horizontal like at the 95% asymptotic threshold, *ℓ*^*∗*^ [56].

**Figure 3:**
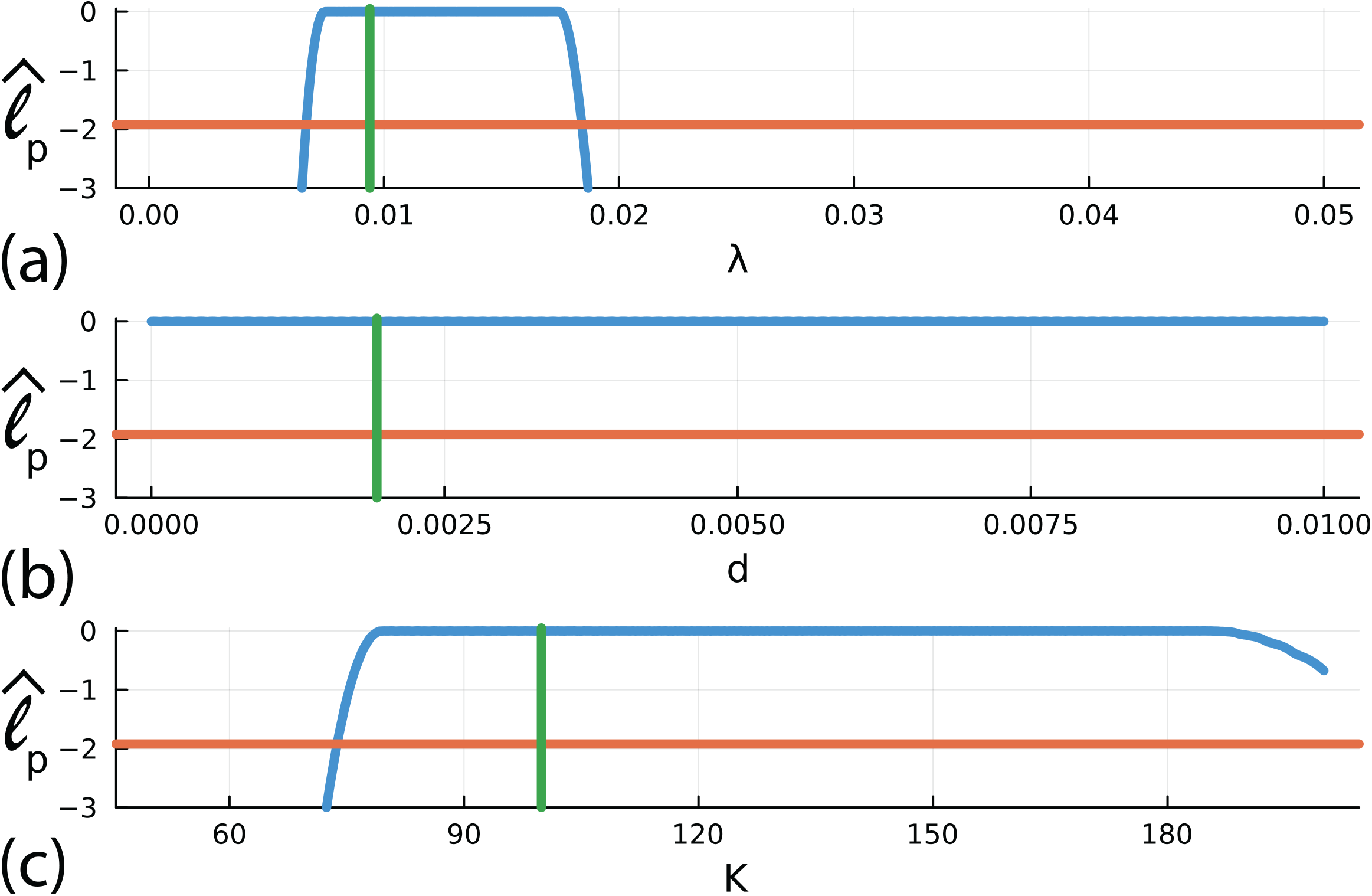
(a)–(c) Univariate profile likelihood functions for *λ, d* and *K*, respectively. Each Profile likelihood function (solid blue) is superimposed with a vertical green line at the MLE and a horizontal red line shoing the asymptotic 95% threshold at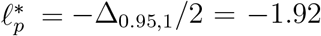, where Δ_*q,n*_ refers to the *q*th quantile of the *χ*^2^ distribution with *n* degrees of freedom [56], here *n* = 1 for univariate profiles. Parameter estimates and 95% confidence intervals are: 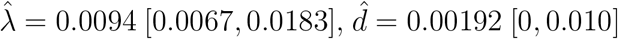 and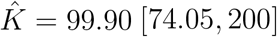.

The univariate profile for *λ* in Figure 3(a) indicates that the noisy data in Figure 2(a) allows us to estimate *λ* reasonably well, at least within the 95% asymptotic confidence interval, 0.0067 *< λ <* 0.0183. In contrast, the relatively flat profiles for *d* and *K* in Figure 3(b)–(c) indicate that the noisy data in Figure 2(a) does not identify these parameters. This is entirely consistent with our observation that this model is structurally non-identifiable. The univariate profile likelihoods indicate there are many choices of *d* and *K* that can be used to match the data equally well. This is a major problem because these flat profiles indicate that incorrect parameter values can match the observations. If, when using mechanistic models, our aim is to link parameter values to biological mechanisms, this means that we can use a model parameterised to reflect an incorrect mechanisms to explain our observations. Here, for example, our data is generated with *K* = 100, but our univariate profile likelihood indicates that we could match the data with *K* = 90 or *K* = 190 which is clearly unsatisfactory if we want to use our data to infer not only the value of *K*, but also to understand and quantify the *mechanism* implied by this parameter value.

Before considering prediction, we first explore the empirical, finite sample coverage properties for our parameter confidence intervals as they are only expected to hold asymptotically [51]. We generate 5000 data realisations in the same way that we generated the single data realisation in Figure 3(a), for the same fixed true parameters. For each realisation we compute the MLE and count the proportion of realisations with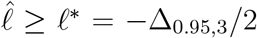, where 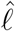 is evaluated at the true parameter values. This condition means the true parameter vector is within the sample-dependent, likelihood-based confidence set. This gives 4900*/*5000 = 98.00% which, unsurprisingly, exceeds the expected 95% asymptotic result due to the fact that the likelihood function is relatively flat. This result is very different to previous implementation of the PWA workflow for identifiable problems where finite sample coverage properties for parameter confidence intervals are very close to the expected asymptotic result [48, 63].

Estimating prediction uncertainties accurately is nontrivial, due to the nonlinear dependence of model characteristics on parameters [71, 72]. The recent PWA workflow addresses identifiability, estimation and prediction in a unified workflow that involves propagating profile-likelihood-based confidence sets for model parameters, to model prediction sets by isolating how different parameters, influences model prediction by taking advantage of the targeting property of the profile likelihood. Previous applications of PWA found that constructing bivariate profile likelihood functions is a useful way to explore relationships between pairs of parameters, and to construct accurate prediction intervals. However, these previous applications have focused on identifiable problems, whereas here we consider the more challenging and realistic question of dealing with non-identifiable and poorly-identifiable models. The idea of using bivariate profile likelihood functions to provide a visual indication of the confidence set of pairs of parameters and the nonlinear dependence of parameter estimates on each other has been long established in the nonlinear regression literature [3]. Here, and in the PWA, we use a series of bivariate profile likelihoods to provide a visual interpretation of the relationship between various pairs of parameters, as well as using these profile likelihood functions to make predictions so that we establish how the variability in parameters within a particular confidence set map to predictions that are more meaningful for collaborators in the life sciences. One of the benefits of working with relatively simple models with a small number of parameters is that we can take the union of predictions from various made using all possible bivariate profile likelihoods and compare these predictions from the full likelihood. Our previous work has compared making such predictions using the union of univariate profile likelihood-based predictions, and union of bivariate profile likelihood-based predictions and predictions from the full likelihood function, funding that working with bivariate profile likelihood-based predictions leads to accurate predictions at a significantly reduced computational overhead when using the full likelihood function.

To construct bivariate profile likelihood functions by setting: (i) *ψ* = (*λ, d*)^⊤^ and *ω* = *K*; (ii) *ψ* = (*λ, K*)^⊤^ and *ω* = *d*; and, (iii) *ψ* = (*d, K*)^⊤^ and *ω* = *λ* and evaluating Equation 10. Again, just like the univariate profile likelihood functions, evaluating 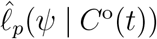 is straightforward because we have already evaluated 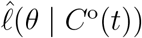 on a 500^3^ regular discretisation of *θ*. For example, to evaluate the bivariate profile likelihood for *ψ* = (*λ, d*)^⊤^ we take each pair of (*λ*_*j*_, *d*_*k*_) values on the discretisation of *θ* and compute the index *l* that maximises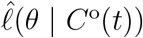, where *θ* is evaluated on the uniform discretisation (*λ*_*j*_, *d*_*k*_, *K*_*l*_) to give the bivariate profile shown in Figure 4(a) where we superimpose a curve at the 95% asymptotic threshold *ℓ*^*∗*^ and the location of the MLE. The shape of the bivariate profile shows that there is a narrow region of parameter space where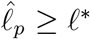, and the shape of this region illustrates how estimates of *λ* and *d* are correlated (i.e. as *λ* increases the value of *d* required to match the data also increases). This narrow region of parameter space where 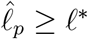 extends right across the truncated region of parameter space considered.

**Figure 4:**
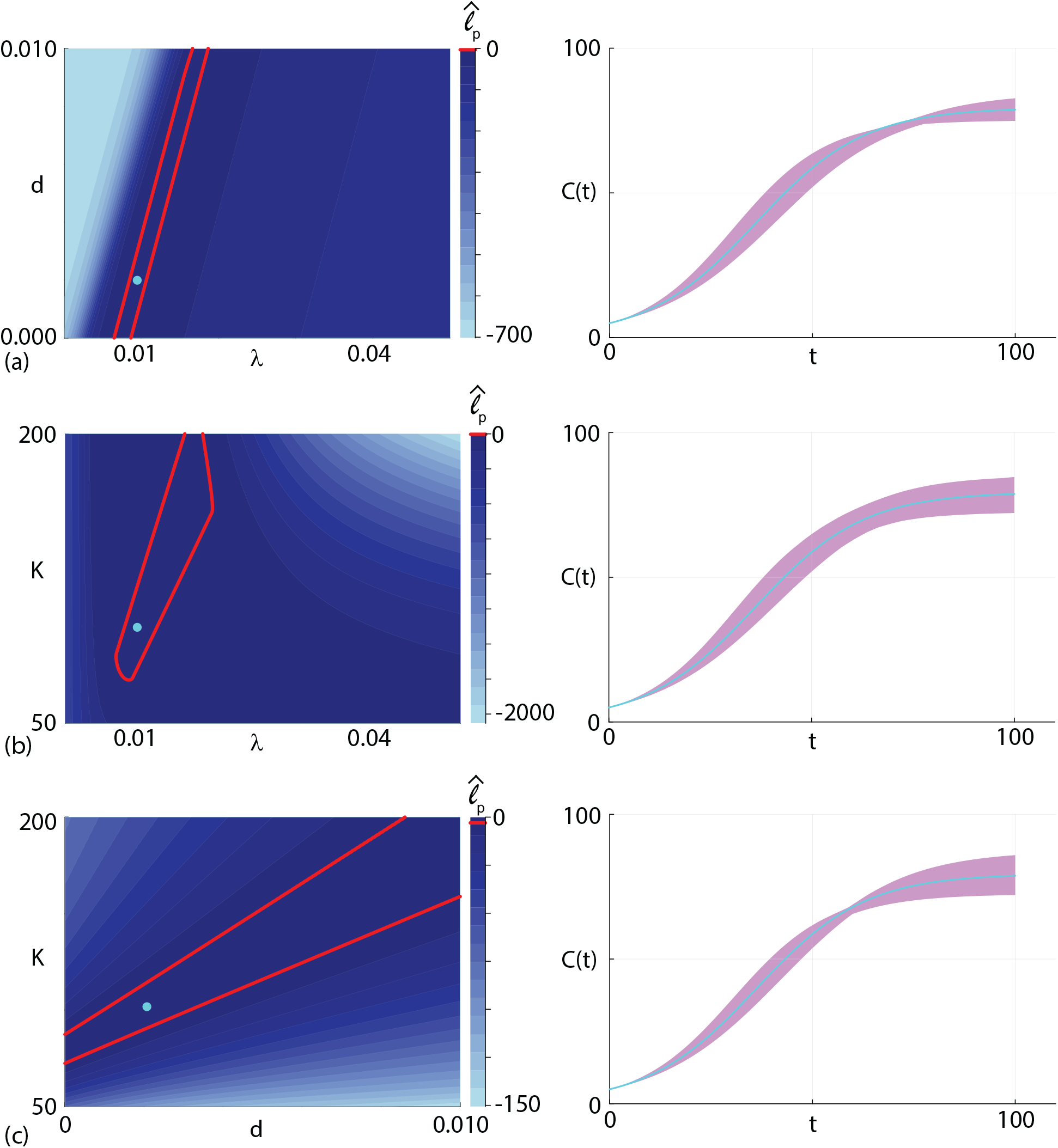
(a)–(c) Bivariate profile likelihood functions for *ψ* = (*λ, d*), *ψ* = (*λ, K*) and *ψ* = (*d, K*), respectively, shown in the left column, together with the associated prediction intervals in the right column. Contour plots in the left column show 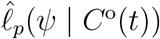 for each choice of *ψ*, as indicated. Each contour plot is superimposed with a contour showing the asymptotic threshold *ℓ*^*∗*^ = *−*Δ_0.95,2_*/*2 = *−*3.00 (solid red curve) and the location of the MLE (cyan disc). Each of the bivariate profiles in the left column is constructed using a 500 *×* 500 uniform mesh, and parameter values lying within the contour where *ℓ ≥ ℓ*^*∗*^ are propagated forward to form the profile–wise prediction intervals in the right column (purple shaded region). Each prediction interval is superimposed with the MLE solution (cyan solid curve).

To understand how the uncertainty in *ψ* = (*λ, d*)^⊤^ impacts the uncertainty in the prediction of the mathematical model, Equation 4, we take those values of (*λ*_*j*_, *d*_*k*_) for which 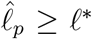 (i.e. those values of *θ* lying within the region enclosed by the threshold, including values of *θ* along the boundary) and solve Equation 4 using the associated value of the nuisance parameter, *K*_*l*_, that was identified when we computed the bivariate profile likelihood. Solving Equation 4 for each choice of (*λ*_*j*_, *d*_*k*_, *K*_*l*_) within the set of parameters defined by the asymptotic threshold provides a family of solutions from which we can compute a curvewise prediction interval, shown in the right most panel of Figure 4(a) where we have also superimposed the MLE solution. This prediction interval explicitly shows us how uncertainty in the values of (*λ, d*)^⊤^ propagates into uncertainty in the model prediction, *C*(*t*).

We now repeat the process of computing the bivariate profile likelihoods for *ψ* = (*λ, K*) and *ψ* = (*d, K*), shown in Figure 4(b)–(c), respectively. Here we see that the bivariate profile likelihood functions provides additional insight beyond the univariate profiles in Figure 3. For example, here we see that bivariate profile likelihood for *ψ* = (*d, K*) is similar to the bivariate profile for *ψ* = (*λ, d*) in the sense that there is a large region, extending right across the parameter space, where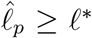. In contrast, for *ψ* = (*λ, K*) we see that the curvature of the bivariate profile likelihood is such that the region where 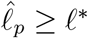 is mostly contained within the region of parameter space considered. The profile wise prediction intervals for each bivariate profile likelihood is given in the right–most panel in Figure 4(a)–(c) together with the MLE solution. Comparing the profile-wise prediction intervals in the Figure 4 illustrates how differences in parameters affect these predictions. For example, the prediction intervals in Figure 4(b)-(c), which explicitly involves bivariate profile likelihoods involving *K* are wider at late time than the prediction interval in Figure 4(a). This reflects the fact that the carrying capacity density, *K* affects the late time solution of Equation 3.

In addition to constructing profile-wise prediction intervals to explore how uncertainty in pairs of parameters impacts model predictions, we can also form an approximate prediction interval by taking the union of the three profile-wise prediction intervals. This union of prediction intervals, shown in Figure 5, compares extremely well with the gold-standard prediction interval obtained using the full likelihood function. In this case it is straightforward to construct the full likelihood prediction interval because we have already evaluated 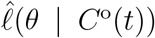 at each of the 500^3^ mesh points, and we simply solve Equation 3 for each value of *θ* for which 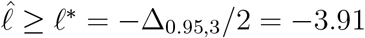 and use this family of solutions to form the curvewise gold-standard prediction interval.

**Figure 5:**
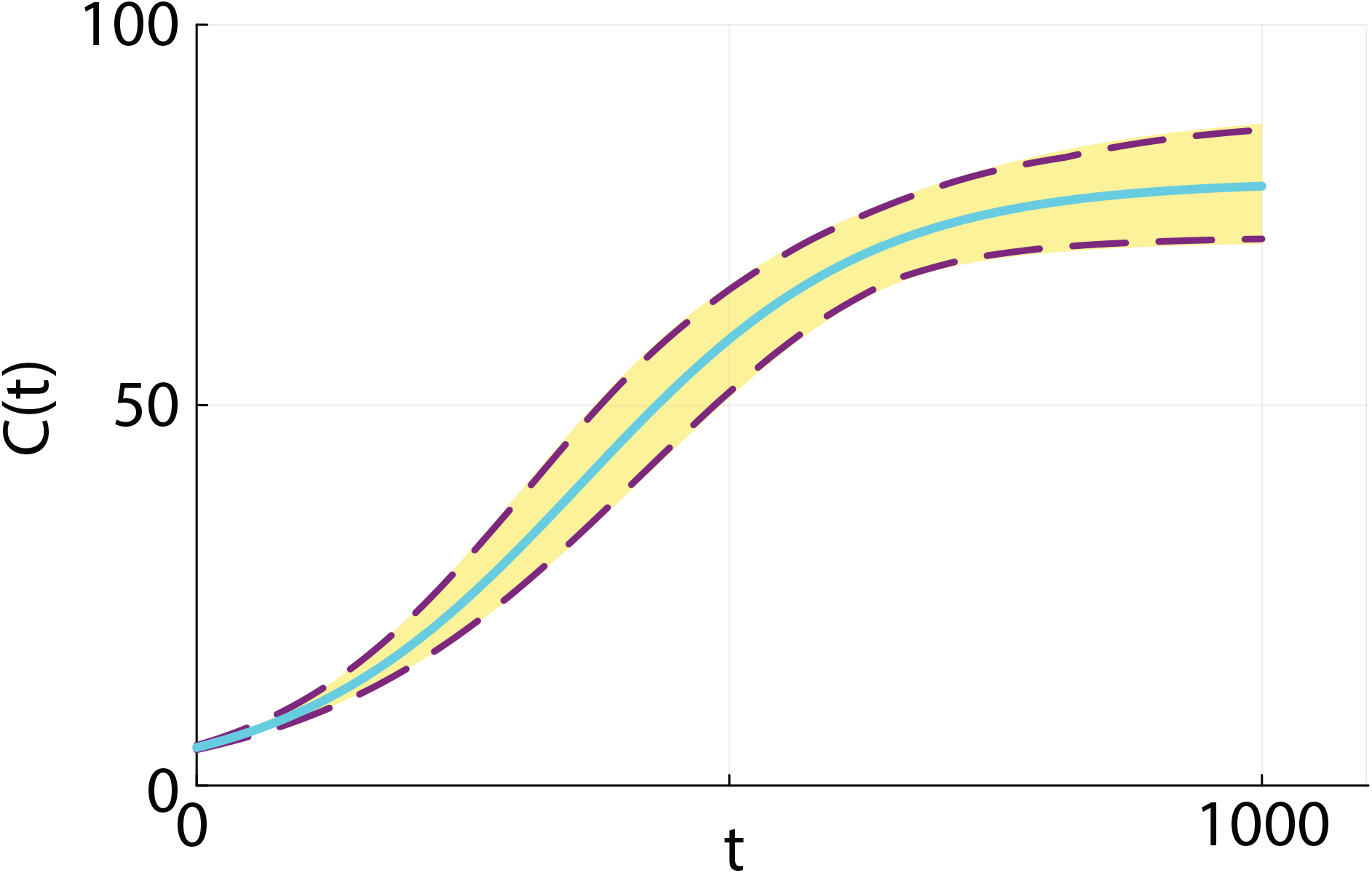
Prediction interval comparison. The prediction interval constructed from the gold standard full likelihood (solid gold region) is superimposed with the MLE solution (cyan solid curve) and the union of the upper and lower profile–wise intervals in Figure 4 (dashed red curves).

Our construction of the approximate full prediction interval is both practically and computationally convenient. This approach is practically advantageous in the sense that building profile-wise prediction intervals like we did in Figure 4 provides insight into how different pairs of parameters impacts model predictions in a way that is not obvious when working with the full likelihood function. This process is computationally advantageous since working with pairs of parameters remains computationally tractable for models with larger number of parameters where either directly discretizing or sampling the full likelihood function becomes computationally prohibitive. Finally, this example indicates that the PWA workflow presented previously for identifiable problems can also be applied to structurally non-identifiable problems. The main difference between applying the PWA workflow for identifiable and structurally non-identifiable problems is that the former problems involve determining regions in parameter space where 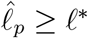 implicitly defined by the curvature of the likelihood function. In contrast, the region of parameter space where 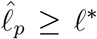 for structurally non-identifiable problems is determined both by the curvature of the likelihood function and the user–defined bounds on the regions of parameter space considered. These bounds can often be imposed naturally, such as requiring that certain parameters be non-negative as in the case of *K >* 0 in Equation 3. Alternatively these bounds can be also be prescribed based on the experience of the analyst, such as our results in Figure 3–4 where we limited our parameter space to the region where 0.0001 *< λ <* 0.05. In this case we chose the interval to be relatively broad, guided by our knowledge that the true value lies within this interval. Later, in Section 2.3 we will deal with a more practical case involving real measurements where we have no *a priori* knowledge of the true parameter values.

The results in this Section for Equation 3 are insightful in the sense that they provide the first demonstration that the PWA workflow can be applied to structurally non-identifiable problems, however we now turn our attention to a class problems that we believe to be of greater importance, namely a practically non-identifiable problem where some parameters are well identified by the data, whereas others are not.

### 2.3 Richards’ model

We now consider estimating parameters in the Richards’ model, Equation 6 to describe the coral reef recovery data in Figure 1(a)–(b). To solve Equation 6 we require values of (*λ, β, K, C*(0))^⊤^, and we assume that the noisy field data is associated with a Gaussian additive noise model with a pre-estimated standard deviation *σ* = 2 [32, 59]. Therefore, unlike the work in Section 2.2 where it was relatively straightforward to discretize the three–dimensional parameter space and calculate the MLE and associated profile likelihood functions simply by searching the discretised parameter space, we now perform all calculations using a standard implementation of the Nelder-Mead numerical optimisation algorithm using simple bound constraints within the NLopt routine [36]. Figure 5(a) shows the data superimposed with the MLE solution of Equation 6 where we have 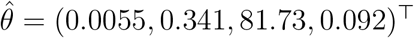 and we see that the shape of the MLE solution provides a very good match to the data. The parameter *λ* has units of [*/*day] and the exponent *β* is dimensionless. Both *K* and *C*(0) measure coral densities in terms of the % of area covered by hard corals.

Before we proceed to consider the curvature of the likelihood function we construct a gold standard full likelihood prediction interval using rejection sampling to find 5000 estimates of *θ* for which *ℓ ≥ ℓ*^*∗*^ = *−*Δ_0.95,4_*/*2 = *−*4.74. Evaluating the solution of the Richards’ model, Equation 7 for each identified *θ* gives us 5000 traces of *C*(*t*), which we use to construct the prediction interval shown by the gold shaded interval in Figure 5(b).

To proceed with the PWA workflow, we first assess the practical identifiability of each parameter by constructing four univariate profile likelihood functions by evaluating Equation 10 with *ψ* = *λ, ψ* = *β, ψ* = *K* and *ψ* = *C*(0), respectively. Profiles are constructed across a uniform mesh of the interest parameter, and the profile likelihood is computed using numerical optimisation. The four univariate profile likelihood functions are given in Figure 7 where each univariate profile likelihood is superimposed with a vertical green line at the MLE and horizontal line at the asymptotic 95% threshold, *ℓ*^*∗*^. The univariate profile for *λ* is flat, meaning that the data is insufficient to precisely identify *λ*. In contrast, profiles for *β, K* and *C*(0) are relatively well–formed about a single maximum value at the MLE. Together, these profiles indicate that we have a commonly encountered situation whereby we are working with a structurally identifiable mathematical model, but the data we have access to does not identify all the parameters. Instead, the data identifies some of the parameters, *β, K* and *C*(0), whereas the data does not contain sufficient information to identify *λ*. This problem is practically non-identifiable, and we will now show that the PWA workflow can be applied to this problem and that this approach yields useful information that is not otherwise obvious.

**Figure 6:**
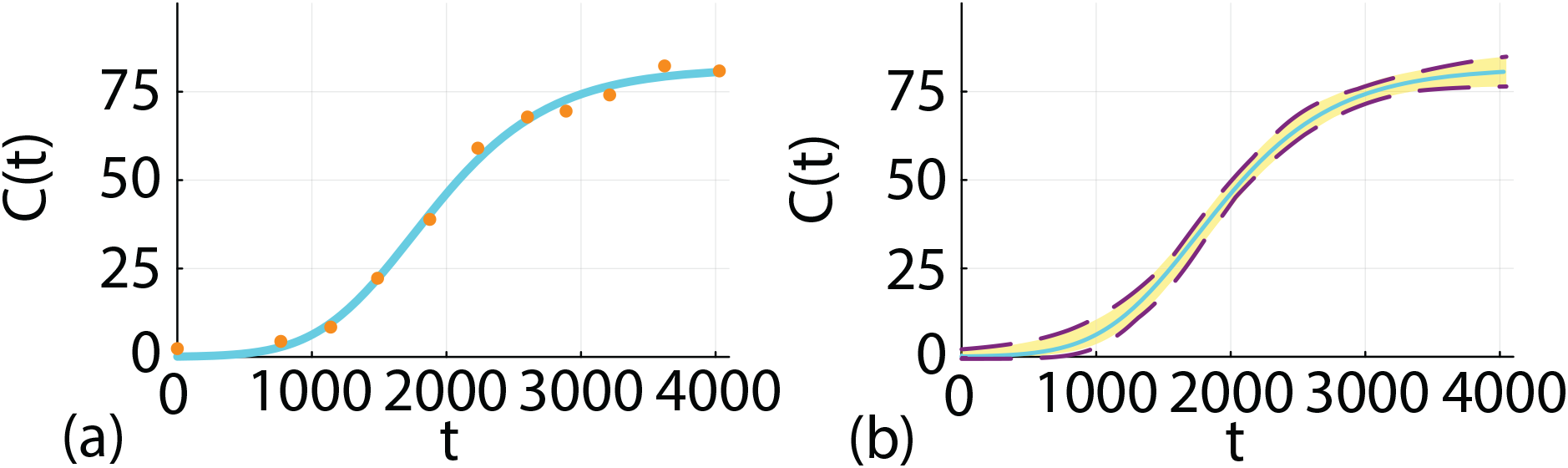
(a) Field data showing the time evolution of the percentage total hard coral cover, *C*°(*t*) (orange discs) after some disturbance at Lady Musgrave Island monitoring site 1 [62]. The MLE solution of the Richards’ model, Equation 6 with 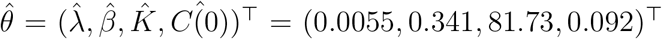 is superimposed (solid cyan curve). (b) Full likelihood-based prediction interval (gold region) superimposed on the MLE solution (solid cyan curve). The full likelihood prediction interval is formed by using rejection sampling to find 1000 values of *θ* with *ℓ ≥ ℓ*^*∗*^ = *−*Δ_0.95,4_*/*2 = *−*4.74, and each value of *θ* is then propagated forward, via Equation 7, to give a family of solution curves from which we construct the prediction interval.

**Figure 7:**
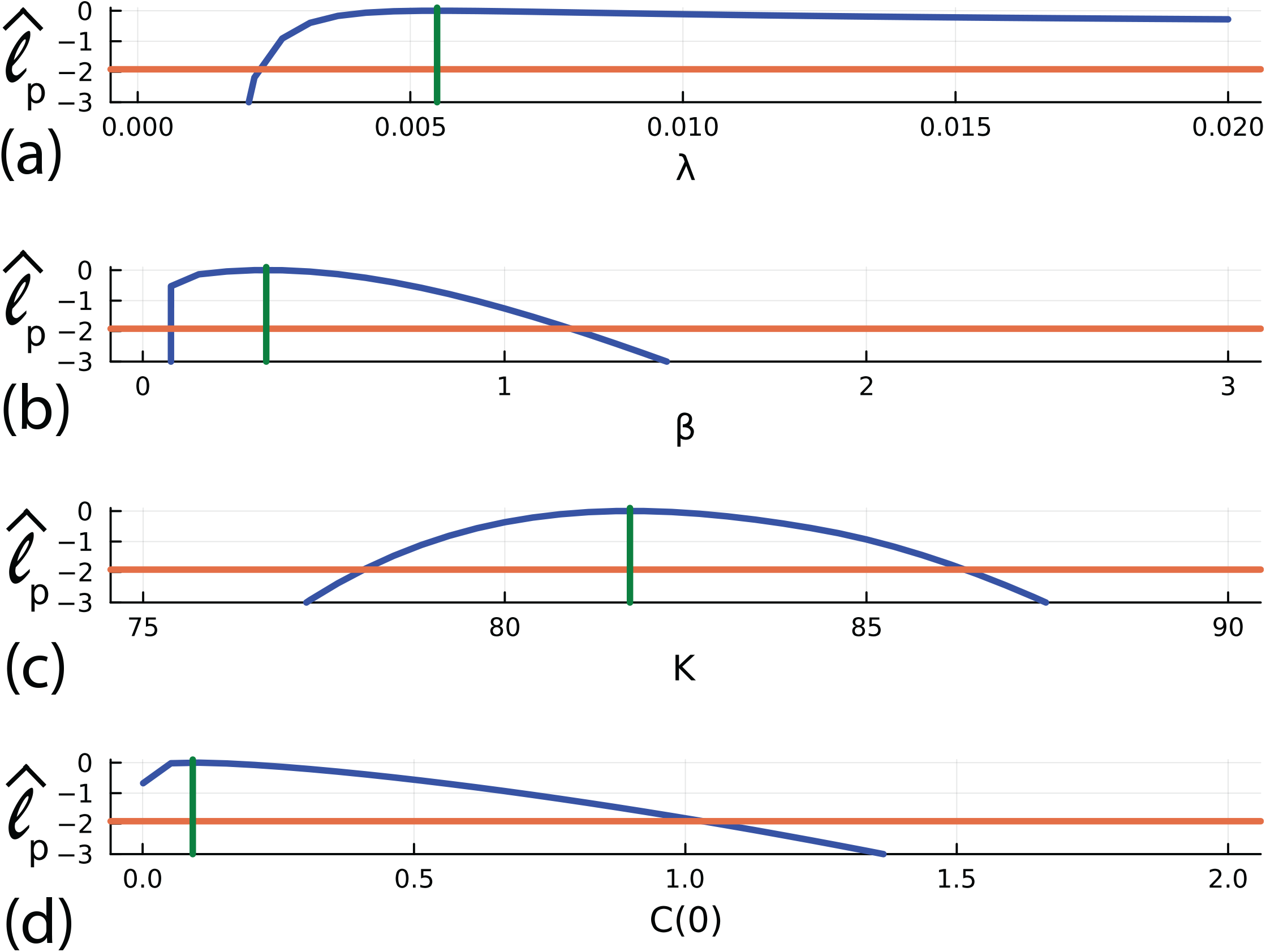
(a)–(d) Univariate profile likelihood functions for *λ, β, K* and *C*(0), respectively. Each Profile likelihood function (solid blue) is superimposed with a vertical green line at the MLE and a horizontal red line shoing the asymptotic 95% threshold at *ℓ*^*∗*^ = *−*Δ_0.95,1_*/*2 = *−*1.92, where Δ_*q,n*_ refers to the *q*th quantile of the χ^2^ distribution with *n* degrees of freedom [56], here *n* = 1 for univariate profiles. Parameter estimates and 95% confidence intervals are: 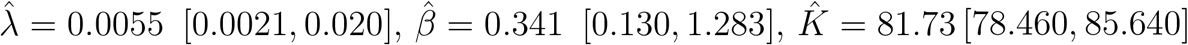 and 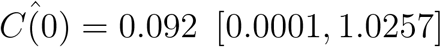.

As with the logistic model with harvesting, before considering prediction we first explore the empirical, finite sample coverage properties. We generate 5000 data realisations in the same way that we generated the single data realisation in Figure 3(a), for the same fixed true parameters. For each realisation we compute the MLE and count the proportion of realisations with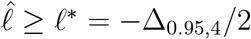, where 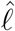 is evaluated at the true parameter values. This gives 4780*/*5000 = 95.60%, which is close to the expected asymptotic result, regardless of the non-identifiability of *λ*.

We now consider prediction by computing various bivariate profile likelihoods. Results in Figure 8(a)–(f) show the six bivariate profile likelihoods:

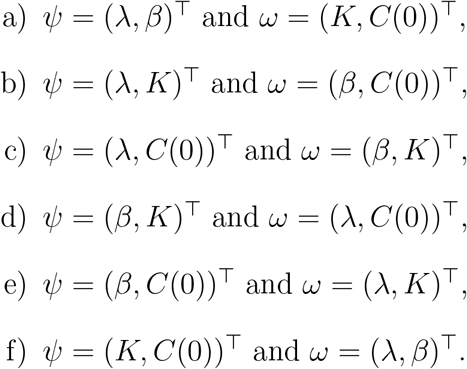

**Figure 8:**
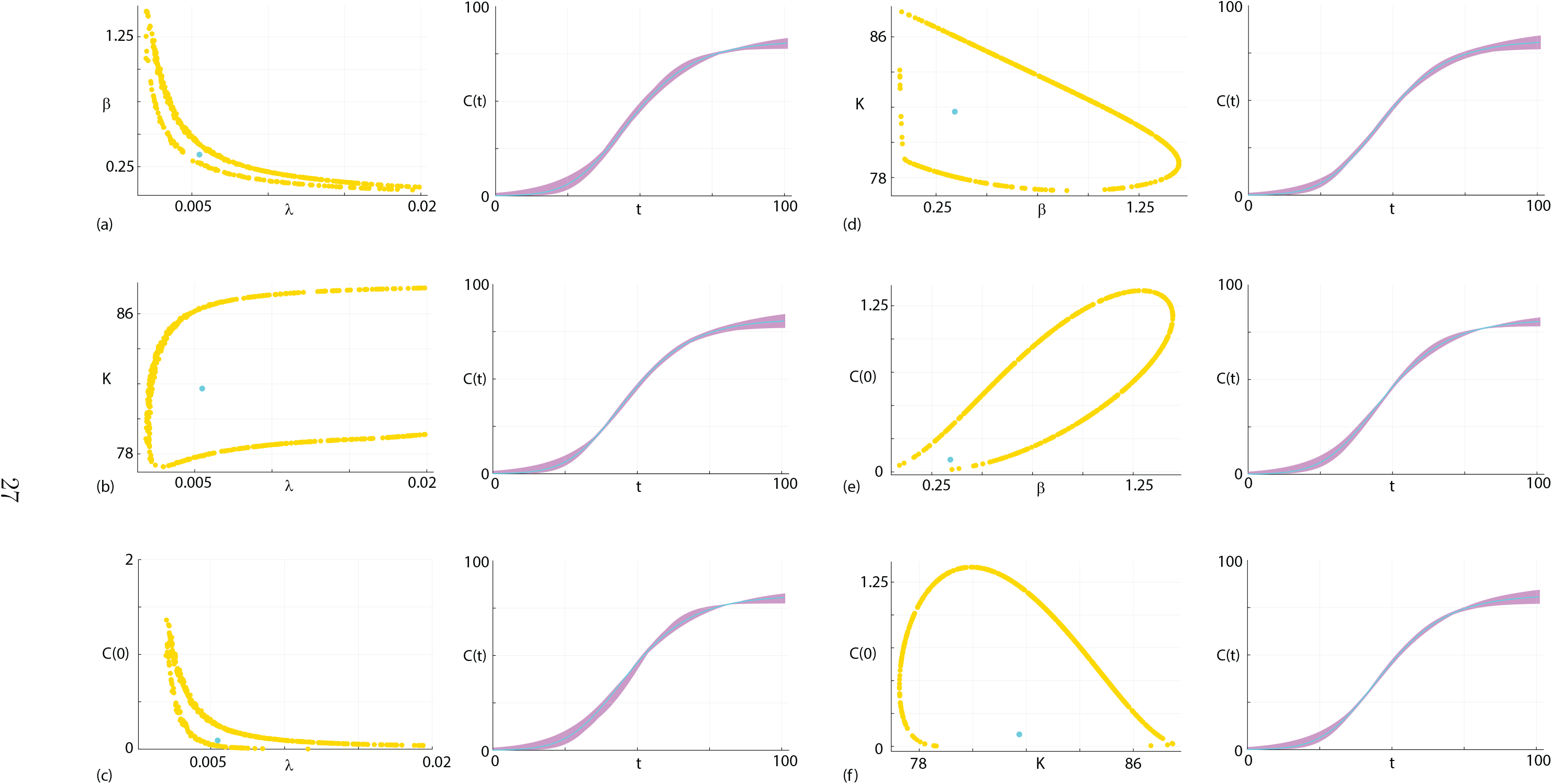
(a)–(f) Approximate contours of six bivariate profile likelihood functions for *ψ* = (*λ, β*), *ψ* = (*λ, K*), *ψ* = (*λ, C*(0)), *ψ* = (*β, K*), *ψ* = (*β, C*(0)) and *ψ* = (*K, C*(0)), respectively, shown in the left of each subfigure. Each bivariate profile is presented adjacent to the associated prediction intervals in the right of each subfigure. Each bivariate profile likelihood is illustrated using 400 randomly identified points along the 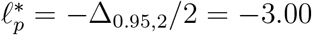 contour (yellow dots) together with the MLE (cyan disc). For each bivariate profile likelihood we propagate each of the 500 values of *θ* along the *ℓ*^*∗*^ = *−*Δ_0.95,2_*/*2 = *−*3.00 forward to give 500 traces of *C*(*t*) that are used to form the profile-wise prediction intervals in the right of each subfigure (purple shaded region). Each prediction interval is superimposed with the MLE solution (cyan solid curve).

Each bivariate profile likelihood is superimposed with the MLE (cyan dot), and following Simpson and Maclaren [63] we randomly identify 500 points along the boundary where *ℓ*^*∗*^ = *−*Δ_0.95,2_*/*2, corresponding to the asymptotic 95% threshold. This approach to propagate confidence sets in parameters to confidence sets in predictions is different to the approach we took in Section 2.3 where we propagated all gridded parameter values for which *ℓ*_*p*_ *≥ ℓ*^*∗*^, but here we focus only on the boundary values where *ℓ*_*p*_ = *ℓ*^*∗*^. As we will later show, this approach of focusing on the boundary values is a computationally convenient way to identify parameter values which, when propagated forward to the model prediction space, forms an accurate prediction interval in terms of comparison with the gold–standard full likelihood approach. Furthermore, the shape of these identified *ℓ*_*p*_ = *ℓ*^*∗*^ contours provides important qualitative insight into parameter identifiability. For example, the contours in Figure 8(a) for the *ψ* = (*λ, β*)^⊤^ bivariate profile is a classic *banana*-shaped profile consistent with our previous observation that *λ* is not identifiable. Similarly, the bivariate profile for *ψ* = (*λ, K*)^⊤^ shows that this contour extends right across the truncated regions of parameter space considered. Again, this is consistent with *λ* not being identified by this data set. In contrast, other bivariate profiles such as *ψ* = (*K, β*)^⊤^ and *ψ* = (*C*(0), *β*)^⊤^ form closed contours around the MLE and are contained within the bounds of the parameter space.

For each bivariate profile in Figure 8 we propagate the 500 boundary values of *θ* forward by solving Equation 6 to give 500 traces of *C*(*t*) from which we can form the profile wise predictions shown in the right–most panel of each subfigure. Propagating the boundary values of *θ* forward allows us to visualise and quantify how variability in pairs of parameters affects the prediction intervals. In this instance we can compare prediction intervals for identifiable pairs of parameters, such as those in Figure 8(d)–(e) with the prediction intervals for pairs of parameters that are not identifiable, such as those in Figure 8(a)–(b). In this instance we see that the width of the prediction intervals dealing with identifiable target parameters appear to be, in general, narrower than the prediction intervals for target parameters that involve non-identifiable parameters.

Taking the union of the profile wise prediction intervals identified in Figure 8 gives us the approximate prediction intervals shown in Figure 6(b) where we see that the union of the profile-wise prediction intervals compares very well with the gold-standard prediction interval formed using the full likelihood approach. While this comparison of prediction intervals in Figure 6(b) indicates that working with the full likelihood and the bivariate profile likelihoods leads to similar outcome interms of the overall prediction interval for *C*(*t*), this simple comparison masks the fact that working with the bivariate profile likelihood functions provides far greater insight by using the targeting property of the profile likelihood function to understand the how the identifiability/non-identifiability of different parameters impacts prediction.

## 3 Conclusion and Outlook

In this work we demonstrate how a systematic likelihood-based workflow for identifiability analysis, parameter estimation, and prediction, called *Profile-Wise Analysis* (PWA), applies to non-identifiable models. Previous applications of the PWA workflow have focused on identifiable problems where profile-wise predictions, based on a series of bivariate profile likelihood functions, provide a mechanistically insightful way to propagate confidence sets from parameter space to profile-wise prediction confidence sets. Taking the union of the profile-wise prediction intervals gives an approximate global prediction interval that compares well with the gold standard full likelihood prediction interval [63]. In addition, empirical, finite sample coverage properties for the PWA workflow compare well with the expected asymptotic result. In this study we apply the same PWA workflow to structurally non-identifiable and practically non-identifiable mathematical models. Working with simple mathematical models allows us to present the PWA in a didactic format, as well as enabling a comparison of the approximate PWA prediction intervals with the gold-standard full likelihood prediction intervals. In summary we find that the PWA workflow approach can be applied to non-identifiable models in the same way that as for identifiable models. For the structurally non-identifiable models the confidence sets in parameter space are partly determined by user-imposed parameter bounds rather than being determined solely by the curvature of the likelihood function. Since the likelihood function is relatively flat we find that finite sample coverage properties exceed the expected asymptotic result. For the practically non-identifiable models we find that some parameters are well–identified by the data whereas other parameters are not. In this case the PWA workflow provides insight by mechanistically linking confidence sets in parameter space with confidence sets in prediction space regardless of the identifiability status of the parameter.

Demonstrating how the PWA workflow applies to non-identifiable mathematical models is particularly important for applications in mathematical biology because non-identifiability is commonly encountered. Previous approaches for dealing with non-identifiability have included introducing model simplifications, such as setting parameter values to certain default values [59, 73], or seeking some combination of parameters that reduces the dimensionality of the parameter space [30,61,70]. One of the challenges with these previous approaches is that they are often administered on a case-by-case basis without providing more general insight. In contrast, the PWA workflow can be applied to study parameter identifiability and to propagate confidence sets in parameter values to prediction confidence sets without needing to find parameter combinations to address non-identifiability, therefore providing a more systematic approach.

There are many options for extending the current study. In this work we have chosen to deal with relatively straightforward mathematical models by working with ODE-based population dynamics models that are standard extensions of the usual logistic growth model. Working with simple, canonical mathematical models allows us to compare our approximate PWA workflow prediction intervals with gold-standard full likelihood results, as well as making the point that parameter non-identifiability can arise even when working with seemingly simple mathematical models. In this work we have applied the PWA to simple models with just three or four parameters, and previous implementations have dealt with models with up to seven parameters [47,62], and future applications of the PWA to more complicated mathematical models with many more unknown parameters is of high interest. Unfortunately, however, under these conditions it becomes computationally infeasible to compare with predictions from the gold-standard full likelihood approach so we have not dealt with more complicated models in the current study. While we have focused on presenting ODE-based models with a standard additive Gaussian measurement model, the PWA approach can be applied to more complicated mathematical models including mechanistic models based on PDEs [19, 59], as well as systems of ODEs and systems of PDEs [48], or more complicated mathematical models that encompass biological heterogeneity, such as multiple growth rates [7]. Here we have chosen to work here with Gaussian additive noise because this is by far the most commonly–used approach to relate noisy measurements with solutions of mathematical models [32], but our approach can be used for other measurement models, such as working with binomial measurement models [43,64] or multiplicative measurement models that are often used to maintain positive predictions [48], as well as correlated noise models [40]. Another point for future consideration is that the present study has dealt with prediction intervals in terms of the mean trajectories. When it comes to considering different measurement models it is also relevant to consider how trajectories for data realisations are impacted, since our previous experience has shown that different measurement models can lead to similar mean trajectories but very different data realisations [64]. In addition, these approaches can also be applied to stochastic models if a surrogate likelihood is available, as discussed in [63] and implemented by Warne et al. [75].

## Data Accessibility

Julia implementations of all computations are available on GitHub https://github.com/ProfMJSimpson/NonidentifiableWorkflow.

## Authors’ Contributions

OJM and MJS conceived the study. MJS carried out the modelling simulations, developed computational algorithms, and wrote code to implement the algorithms, and interpreted the results. OJM and MJS drafted the manuscript, and approved the final version.

## Acknowledgements

We thank two referees for helpful suggestions.

